# TrAEL-seq captures DNA replication dynamics in mammalian cells

**DOI:** 10.1101/2025.06.09.658559

**Authors:** Neesha Kara, Laura Biggins, Vera Grinkevich, Alex Whale, Paola Garran-Garcia, Jhanavi Srinivasan, Peter J. Rugg-Gunn, Simon Andrews, Aled Parry, Helen M. R. Robinson, Jonathan Houseley

## Abstract

Precise DNA replication is critical to the maintenance of genome stability, and the DNA replication machinery is a focal point of many current and upcoming chemotherapeutics. TrAEL-seq is a robust method for profiling DNA replication genome-wide that works in unsynchronised cells and does not require treatment with drugs or nucleotide analogues. Here, we provide an updated method for TrAEL-seq including multiplexing of up to 6 samples that dramatically improves sample quality and throughput, and we validate TrAEL-seq in multiple mammalian cell lines. The updated protocol is straightforward and robust yet provides excellent resolution comparable to OK-seq in mammalian cell samples. High resolution replication profiles can be obtained across large panels of samples and in dynamic systems, for example during the progressive onset of oncogene induced senescence. In addition to mapping zones where replication initiates and terminates, TrAEL-seq is sensitive to replication fork speed, revealing effects of both transcription and proximity to replication Initiation Zones on fork progression. Although forks move more slowly through transcribed regions, this does not have a significant impact on the broader dynamics of replication fork progression, which is dominated by rapid fork movement in long replication regions (>1Mb). Short and long replication regions are not intrinsically different, and instead replication forks accelerate across the first ∼1 Mb of travel such that forks progress faster in the middle of regions lying between widely spaced Initiation Zones. We propose that this is a natural consequence of fewer replication forks being active later in S phase when these distal regions are replicated and there being less competition for replication factors.

## Introduction

DNA replication is initiated by the firing of replication origins. Origins in budding yeast are exactly defined by specific DNA sequences but in mammalian cells are heterogeneously distributed and poorly defined with replication initiation events concentrated in, though not restricted to, large Initiation Zones of up to 150kb [1-4]. The pattern of Initiation Zone usage varies between cell types, but is robust across multiple cells in a population, showing that Initiation Zone usage is specified even if individual initiation sites are not [1, 5, 6].

Discovering the set of Initiation Zones used in any given cell-type is a substantial undertaking, and we therefore have little understanding of how active Initiation Zones are specified in different cells, or how variable Initiation Zone usage really is.

Replication initiation can be profiled genome-wide by isolation of replication intermediates (OK-seq, SNS-seq, Bubble-seq) though these methods involve significant biochemical purifications [1, 2, 7]. Flow sorting of precise S-phase fractions identifies early origins though spatial resolution is lower (Repli-seq)[8, 9], while nucleotide analogue incorporation reveals initiation events if cells are synchronised or long read single molecule sequencing applied (INI-seq, DNAscent) [3, 10]. DNA end mapping methods also yield replication profiles: GLOE-seq, which detects all single-stranded DNA (ssDNA) ends can profile Okazaki fragments in a DNA ligase mutant [11], while TrAEL-seq, which detects free single stranded DNA 3’ ends in double stranded DNA has a strong selectivity for leading strand ends and provides DNA replication maps even on unsynchronised wildtype cells [12]. Finally, since DNA polymerase usage differs between leading and lagging strands, replication profiles can be constructed using polymerase mutants that incorporate excessive ribonucleotides on one strand or the other (PU-seq) [13, 14]. There is considerable discordance between these methods in the fine scale mapping of initiation events, but the general clustering events into Initiation Zones is widely reproduced [15] DNA replication forks traverse large replication zones, often >1 Mb in mammalian cells, with particularly large replication zones being prone to incomplete replication resulting in fragile site expression if fork progression is impaired [1, 16, 17]. Replication fork speed is regulated to ensure complete, high fidelity replication, being increased in large replication zones where replication continues late in S phase [18-20], but reduced under conditions of high oxidative damage or transcriptional load to minimise the hindrance to fork progression from obstacles [21, 22]. Although DNA fibre assays are widely used to measure replication fork processivity, these assays do not provide genomic information and therefore the effects of genome structure and localised obstacles cannot be determined. TrAEL-seq has great potential in this regard as it can detect the distribution of replication forks across the genome in both asynchronous and synchronised populations [23, 24].

Impediments to DNA replication cause replication stress. Endogenous sources of replication stress include nucleotide depletion, unrepaired DNA lesions, misincorporated ribonucleotides, repetitive DNA sequences, secondary DNA structures and collisions with transcription units (reviewed in [25]). The mutagenicity of transcription-replication conflicts has been particularly studied and arises both from direct interactions between the replication and transcription machinery, and from the replication inhibitory effects of R-loops [26-28].

Furthermore, exclusion of replication initiation from very large genes creates extended regions requiring contiguous replication; these are prone to under-replication and form fragile sites [16, 29, 30]. However, such mutagenic interactions are necessarily rare, indeed mammalian cells regulate transcription to minimise such transcription-replication conflicts [31], and it is not clear whether the transcriptional machinery routinely delays the passage of replication forks [32].

DNA replication is the target of both classic and targeted chemotherapeutics, and although we have an excellent understanding of the biochemical impact of these treatments, revealing how replication fork processivity is affected and how this is affected by features such as chromatin structure and transcription units remains extremely challenging. We previously demonstrated the utility of TrAEL-seq in profiling DNA replication in unsynchronised steady state cell populations, and here we present a six-way multiplexing modification that dramatically improves quality and throughput while reducing cost. We validate TrAEL-seq in multiple human cell lines and show that TrAEL-seq reveals dynamic changes in replication fork processivity across the genome.

## Results and discussion

### Multiplexed TrAEL-seq in mammalian cells

The throughput of TrAEL-seq library synthesis can be increased by adding a 4 nucleotide in-line barcode to the first TrAEL-seq adaptor allowing libraries to be pooled for processing after initial adaptor ligation (Figure 1A). Although we previously multiplexed pairs of samples [12], the resulting barcode efficiencies were poor so we redesigned the adaptors and changed manufacturer, resulting in dramatically better barcode fidelity and minimal unassigned reads. Of 9 barcodes tested, 6 showed equivalent, high TrAEL-seq efficiency, providing similar read numbers when ligated competitively to a single sample of DLD-1 cells (Figure 1B,S1A), and Replication Fork Directionality (RFD) profiles obtained through this 6-way multiplexing procedure were indistinguishable (Figure 1C).

**Figure 1:**
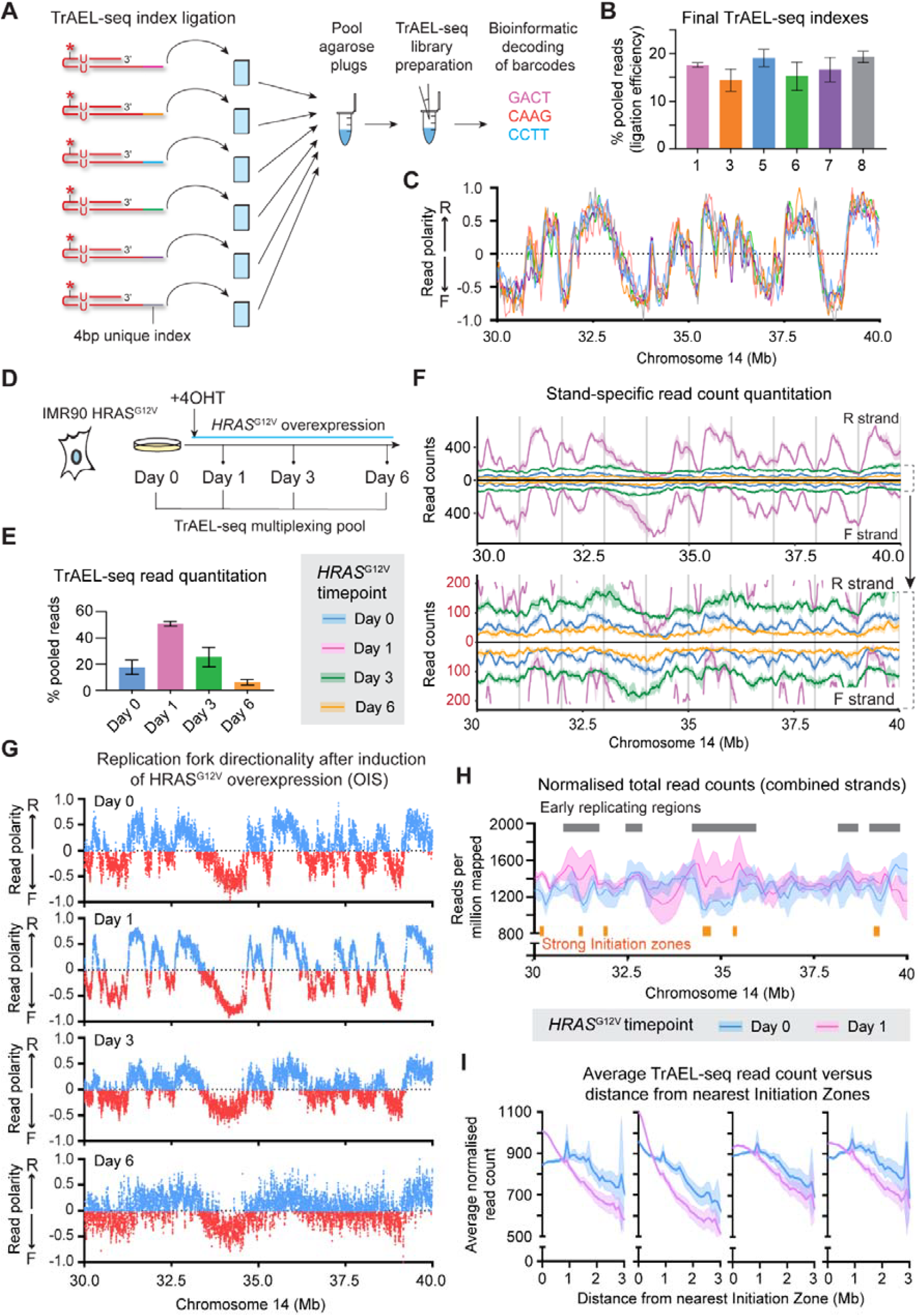
Multiplexed Tr AEL-seq for replication profiling in complex systems. A: Schematic of multiplexed TrAEL-seq – barcoded adaptors are incorporated early in protocol allowing samples to be pooled for processing. Multiple pools can be generated over a period of weeks then all processed into libraries together. B: Read counts from TrAEL-seq adaptors with 6 different indexes ligated competitively to one DNA sample of DLD-1 cells in a single reaction. Some barcodes affect ligation efficiency as the barcode is placed at the ligation junction, but these barcodes show equivalent ligation efficiencies. Error bars ± 1 S.D. n=4. C: Replication fork direction (RFD) plots of an example chromosomal region obtained from the competitive ligation experiment in B. RFD is calculated from read polarity as (R-F)/(R+F), such that positive values represent forks moving left->right and negative values represent forks moving right->left. Calculated in 20kb windows. D: Schematic of oncogene induced senescence timecourse in IMR90 with tetracyclin inducible HRAS^G12V^. E: Number of reads obtained from TrAEL-seq libraries at each time point. Each replicate time course was multiplexed in a separate TrAEL-seq library, with indexes rotated to avoid index-specific effects. Error bars ± 1 S.D. n=4. F: Quantitative profiles of TrAEL-seq reads over an example chromosomal region at different timepoints leading to oncogene induced senescence. Reverse and Forward reads (which capture replication forks moving left->right and right->left respectively) are shown separately. Solid line shows mean signal, shaded band ± 1 S.D. n=4. G: RFD plots across the same example chromosome region shown in F, separated by oncogene induced senescence timepoint. Calculated in 10 kb windows spaced every 1 kb. H: Total TrAEL-seq read count across an example chromosomal region for Day 0 and Day 1 of oncogene induced senescence timecourse. Solid line shows mean signal, shaded band ± 1 S.D. n=4. I: Plots of total TrAEL-seq at increasing distance from replication Initiation Zones. Each plot shows a single biological replicate, solid line shows mean value, shaded band 95% confidence interval.

We rebuilt the TrAEL-seq bioinformatic pipeline to separate reads by inline barcode prior to mapping and introduced a new copy number-aware UMI deduplication step. We further added a quality control routine to ensure TrAEL-seq reads stem from DNA as in theory TrAEL-seq can also detect RNA; although we have observed no evidence of read contamination from abundant RNA species, TrAEL-seq may detect RNA that is complexed with DNA. TrAEL involves tailing of DNA 3’ ends with ATP following by adaptor ligation using an RNA ligase, and could therefore detect any nucleic acids terminating with a 3’ RNA nucleotide through terminal transferase-independent ligation of adaptor to such ends. DNA-derived reads have an invariant T at the first nucleotide from the A-tailing, whereas RNA-derived reads could start with any nucleotide, so by splitting reads into those which start with a T and those which don’t, unexpected sites of read accumulation can be assigned to DNA or RNA depending on representation in the no-T fraction. No-T reads constitute up to 5% of the mapped reads in some libraries, and although these largely follow the replication pattern, it is possible that other features such as nascent transcripts and R-loops contribute at some sites.

TrAEL-seq libraries generated using the refined multiplex protocol are generally of much higher quality and yield more unique reads than libraries generated using the original TrAEL-seq protocol, allowing high resolution analysis (Figure S1B). The improvement in quality was unexpected but is consistent and presumably stems from the re-designed adaptors as these form the only substantive difference in the updated protocol. Reproducibility and specificity are good as RFD plots generated from unsynchronised mammalian cell samples show very similar patterns of replication fork directionality and Initiation Zone s between biological replicates, but clear differences in Initiation Zone usage between lines as expected (Figure S1C). Replication profiles derived by TrAEL-seq also closely match those generated by OK-seq, as evident in a comparison of published TrAEL-seq and OK-seq data generated from activated mouse B cells in different laboratories (Figure S1D). Note that the directionality of replication forks measured by TrAEL-seq and OK-seq is the same as TrAEL-seq detects the 3’ end of the leading strand whereas OK-seq detects the 5’ end of Okazaki fragments [33, 34].

Overall, the revised multiplexed TrAEL-seq protocol increases the quality, throughput and specificity of TrAEL-seq while maintaining the capability of the original method to assay complex systems without synchronisation, sorting, genetic modification or labelling. To aid in setting up the method, we provide a detailed protocol including solutions to common TrAEL-seq problems (Supplemental File 1). Furthermore, ‘How to’ videos for making and handling agarose plugs, as well as the latest protocol version are available at https://www.babraham.ac.uk/our-research/epigenetics/jon-houseley/protocols.

### Replication fork redistribution during oncogene induced senescence

We applied multiplexed TrAEL-seq to profile the dynamic transitions in DNA replication during the onset of oncogene induced senescence (OIS). Expression of HRAS^G12V^ in human IMR90 fibroblasts initially causes hyper-replication (day 1), but this is unsustainable and proliferation soon declines (day 3), entirely ceasing by day 6 as cells enter senescence (Figure 1D)[35, 36]. We acquired 4 replicate OIS time courses (time points at days 0, 1, 3 and 6), with individual libraries in each multiplexed replicate set showing expected differences in read count given the varying proliferation state of the cells over time (Figure 1E): TrAEL-seq reads increased dramatically from Day 0 to Day 1 but were very rare by Day 6 (Figure 1F). Pooling the replicate sets yielded high resolution RFD plots particularly at Day 1, with progressively decreasing resolution to Day 6 as expected since very few cells are replicating by this time (Figure 1G), but to our surprise we did not observe any change in Initiation Zone usage between Day 0 and Day 1 despite the dramatic increase in replication (Figure 1G).

However, read count distribution across chromosomes was altered as hyper replication commenced. At Day 0, TrAEL-seq reads were evenly distributed, whereas at Day 1 reads were noticeably enriched at many early replicating regions (Figure 1H). The strongest Initiation Zones are found in early replicating regions, and peaks of TrAEL-seq reads were often coincident with strong Initiation Zones at Day 1. To quantify this effect, we calculated the average read count in genomic regions as a function of distance from the nearest Initiation Zone, and observed in all four replicate sets that the normalised read count on Day 1 was greater within ∼0.6Mb of Initiation Zones (p=0.005 by t test of AUCs) but relatively depleted at regions >1Mb from Initiation Zones (p=0.049 by t test of AUCs), which are late replicating (Figure 1I). The strength of this effect was variable between biological replicates, but this is to be expected considering the dynamic changes in proliferation both before and after Day 1.

Multiplexed TrAEL-seq can therefore profile DNA replication in complex, dynamic systems to reveal both Initiation Zone usage and replication fork distribution. TrAEL-seq read density varies with replication state and is almost absent in non-proliferating Day 6 cells (Figure 1E,F), and furthermore TrAEL-seq read orientation is highly polarised around Initiation Zones at both Day 0 and Day 1 (Figure 1G). Therefore, the vast majority of TrAEL-seq reads most likely arise from replication forks rather than noise or some other biological process, and although high levels of fork cleavage soon after initiation could give rise to an accumulation of polarised reads around Initiation Zones this is unlikely given the high proliferation rate of the IMR90 cells at Day 1. We therefore attribute this accumulation of TrAEL-seq reads around Initiation Zones to an uneven distribution of replication forks across the genome.

### Variation in replication fork processivity over the human genome

As the change in TrAEL-seq read distribution occurred in response to an oncogenic stimulus, we examined datasets for other transformed lines and observed a read accumulation around Initiation Zones to a greater or lesser extent for all tested including DLD-1, IMR32, NCI-H23, HCT116 and KM-12 (Figure 2A). However, this phenomenon is not unique to transformed cells or in fact to human cells as we also observed increased TrAEL-seq read density near Initiation Zones in human embryonic stem cells (hESCs), in mouse embryonic stem cells and in activated mouse splenic B cells (Figure 2A).

**Figure 2:**
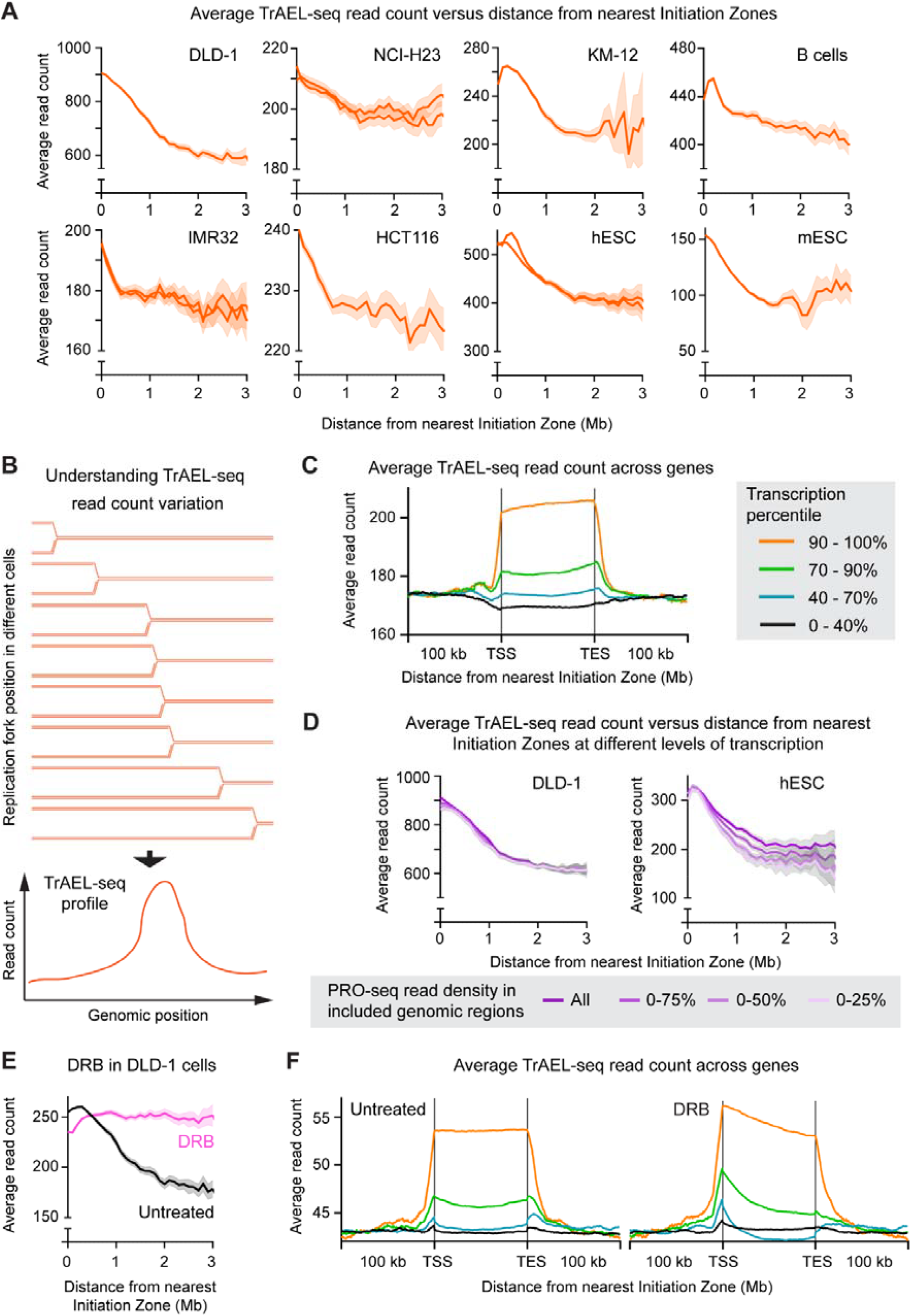
Tr AEL-seq profiling of variable replication fork density. A: Plots of total TrAEL-seq read count at increasing distance from replication Initiation Zones. Each plot shows a different cell line, multiple lines indicate multiple biological replicates where available, solid line shows mean value, shaded band 95% confidence interval. B cell sample shows splenic B cells from C57BL/6 mice activated in culture. Read counts were summed in 50kb windows spaced every 50kb, with regions of altered copy number or aberrant read count removed. Distance was calculated from Initiation Zones determined using OKSeqHMM [45], and average read count determined in 100kb windows of distance from the centre of initation zones. Published datasets are from [23, 34, 46, 47], note that IMR32 and NCI-H23 data were generated prior to the implementation of multiplexed TrAEL-seq, and are therefore derived from lower resolution datasets. B: Schematic showing how TrAEL-seq read density variations arise from variations in replication fork speed averaged across a population. C: Metaplot of TrAEL-seq read count in DLD-1 cells averaged across genes ±100 kb. Genes are stratified for transcriptional activity based on PRO-seq into 0-40%, 40-70%, 70-90% and 90-100% categories. D: Plots of total TrAEL-seq read count at increasing distance from replication Initiation Zones, stratified for nascent transcription level by PRO-seq. Analysis was performed as in A, but the genomic windows included were filtered to remove the top 25%, 50% or 75% of regions based on PRO-seq read count. E: Total TrAEL-seq read count at increasing distance from replication Initiation Zones in untreated DLD-1 cells or cells treated for 2 hours with 100 *µ*M DRB. F: Metaplot of TrAEL-seq read count in DLD-1 cells ± DRB (datasets as in E), averaged across genes ±100 kb as in C.

Importantly, all of these samples derive from unsynchronised, unsorted cell populations assayed at steady state proliferation. Every part of the genome must replicate once per cell cycle, so in populations of unsynchronised cells the density of replication forks should be uniform across the genome as long as replisomes travel at a uniform speed. The fact that more replication forks are detected in some regions of the human genome indicates that replication forks spend longer replicating these regions than other regions of equivalent size. In other words, higher TrAEL-seq read density indicates a genomic region in which replication forks are either pausing more frequently or moving more slowly (Figure 2B).

Decreasing TrAEL-seq density with increasing distance from Initiation Zones therefore suggests that replication forks move faster in regions far from Initiation Zones that are replicated late in S phase, which has been independently observed using single molecule approaches [18, 19].

To confirm by orthogonal methods that higher read counts around Initiation Zones are not a TrAEL-seq artefact, we first performed SSB-seq on DLD-1 cells, which should detect replication forks by extending and labelling both leading and lagging strand ends [37]: in accord with the TrAEL-seq data, SSB-seq reads are also biased towards Initiation Zones (Figure S2A). OK-seq is generally considered the gold standard replication fork profiling method, and we found that OK-seq reads in published B cell data are also biased towards Initiation Zones (Figure S2B)[33]. OK-seq detects Okazaki fragments rather than leading strand ends, but slower Okazaki fragment synthesis would cause Okazaki fragments to remain shorter than the 200bp size cutoff used in OK-seq for longer, and hence become over-represented compared to OK-seq reads from genomic regions in which synthesis is faster.

Many studies have suggested that interactions with the transcription machinery can impair fork processivity and, based on published DLD-1 PRO-seq data, we find that TrAEL-seq reads are enriched ∼20% in the top 10% of highly expressed genes compared to surrounding sequence indicating that replication forks move more slowly through highly transcribed genes (Figure 2C)[38]. Reads lacking an initial T, which are predicted to be RNA-derived are slightly more enriched presumably through detection of nascent transcripts, but the contribution of these to the overall enrichment is <10% (Figure S2C). Furthermore, an equivalent signal was observed when considering reads orientated forward or reverse to the direction of transcription, of which only the reverse reads could conceivably arise from detection of the nascent RNA (Figure S2D). The enrichment is more pronounced for co-directional than head-on interactions, and we saw no convincing evidence of an enrichment at the 3’ end of genes in the head-on interactions that would indicate frequent collisions or break formation (Figure S2E).

Gene density is highest close to Initiation Zones in the human genome, so interactions with transcription may explain the difference in replication fork processivity around Initiation Zones. To investigate this, we generated a high resolution DLD-1 TrAEL-seq dataset by pooling technical replicates to obtain 49 million reads after deduplication, ∼4x our normal sequencing depth (10-20 million after deduplication). We then repeated the analysis of TrAEL-seq read density versus distance from Initiation Zone, excluding regions that were in the top quartile for transcription based on PRO-seq read density (Figure 2D)[38]. This had almost no effect on the profile, so we further excluded probes overlapping the next two quartiles; importantly, even when only including regions in the bottom quartile for transcription (where transcription is essentially undetectable by PRO-seq) the TrAEL-seq read density profile was identical to the complete TrAEL-seq dataset (Figure 2D). Furthermore, an equivalent analysis in hESC showed that exclusion of transcribed regions increased the average TrAEL-seq read enrichment around Initiation Zones (Figure 2D)[39].

We conclude that the variation in replication fork speed with distance from Initiation Zone measured by TrAEL-seq and other methods cannot be caused by interactions with transcription.

This observation contrasts with a recent study reporting that treatment with the transcription inhibitor DRB accelerates replication fork speed [19]. Consistent with that finding, treatment of DLD-1 cells with DRB prevented the reduction in TrAEL-seq read density with distance from Initiation Zones, indicating that fork speed became uniform across the genome (Figure 2E). However, the over-representation of TrAEL-seq reads in highly expressed genes was unaffected except for a slight shift towards the transcriptional start site, which is to be expected given that DRB prevents RNA polymerase II moving from initiating to elongating mode (Figure 2F). This short DRB treatment (2 hours) therefore has only a minimal effect on interactions between replication forks and transcription units detectable by TrAEL-seq, but a huge effect on replication fork progression. This contradiction is most easily resolved by considering that the effects of DRB on replication fork speed stem not from inhibition of transcription but from inhibition of a different kinase, for example casein kinase 2 is inhibited by DRB [40, 41].

Overall, TrAEL-seq provides a wealth of information on replication fork dynamics with high sample throughput and low cost – the method involves no complicated procedures and has proved very reliable in our hands. Importantly, the capability of TrAEL-seq to analyse unsynchronised, unlabelled cell populations both minimises cell culture requirements and allows complex systems to be studied. Here we have leveraged large and varied datasets to assess the effect of transcription on replication forks during normal proliferation: we find that replication fork processivity increases in regions further from Initiation Zones that would be replicated later in S phase, and that transcription locally reduces replication fork processivity. However, these two observations are unconnected and the effect of distance from Initiation Zone (or time in S phase) on replication fork progression rate is not the result of interactions with transcription. It has long been known that replication fork speed is inversely proportional to the number of replication origins that fire, which can be attributed to limiting availability of replication factors (including but not limited to dNTPs) [20, 42-44], and we suggest that replication fork speed increases during S phase simply because inter-origin distances are mostly small (<1Mb) and therefore most forks terminate early in S-phase. Few replication forks traverse regions of >1Mb, and as these regions far from replication Initiation Zones are replicated late in S phase when few other replication forks are active, competition for replication factors is low and the forks can proceed faster.

## Materials and methods

### Cell culture

Undifferentiated H9 hESCs were maintained on Vitronectin-coated plates (Thermo Fisher Scientific, A14700) in TeSR-E8 media (StemCell Technologies, 05990) and Essential 8 Medium (Gibco, A1517001) under 5% O_2_, 5% CO_2_ at 37 °C. DLD-1 cells were maintained in 75 cm^2^ flasks (Thermo Fisher Scientific, 156499 & 159910) in RPMI-1640 media (PAN Biotech, P04-18500) with 10% (v/v) FBS (Fetal Bovine Serum) (Thermo Fisher Scientific, 10270-106), and 100 I.U/mL penicillin-streptomycin (Thermo Fisher Scientific, 15140122) under 5% CO_2_ at 37 °C. Cells were passaged with trypsin-EDTA 0.05% solution (Invitrogen, 25300062) and kept for a maximum of 20 passages. IMR90 ER:HRAS^G12V^ (IMR90 ER:RAS) fibroblasts were gifted from Prof. Masashi Narita at the CRUK Cambridge Institute. Cells were cultured in DMEM (Thermo Fisher Scientific, 31053028) with 10% (v/v) FBS (Thermo

Fisher Scientific, 10270-106), 2 mM L-Glutamine (Thermo Fisher Scientific, 25030081), 100 I.U/mL penicillin-streptomycin (Thermo Fisher Scientific, 15140122) and 1 mM sodium pyruvate (Thermo Fisher Scientific, 11360070) under 5% CO_2_ at 37 °C. For passaging and collection, cells were detached with TrypLE™ Express Enzyme (Thermo Fisher Scientific, 12604013) and centrifuged at 300 g for 3 min. For time course experiments, IMR90 ER:RAS cells were seeded in 10-15 cm dishes (Thermo Fisher Scientific, 150350) and the following day were treated with 4-hydroxy tamoxifen (4OHT) (Sigma, H7904) at a final concentration of 100 nM. 4OHT was replenished every 2-3 days. Cells were fully senescent following 6 days of treatment with 4OHT.

### TrAEL-seq adaptor preparation

Multiplexing variants of TrAEL-seq adaptor 1 were synthesised and PAGE purified by Integrated DNA Technologies (IDT). Sequences for TrAEL-seq adaptors are:

TrAEL-seq adaptor 1 multiplexing index 1:

/5Phos/GACTNNNNNNNNAGATCGGAAGAGCGTCGTGTAGGGAAAGAGTGUAGC A/iBiodT/TGCUACACTCTTTCCCTACACGACGCTCTTCCG

TrAEL-seq adaptor 1 multiplexing index 3:

/5Phos/CAAGNNNNNNNNAGATCGGAAGAGCGTCGTGTAGGGAAAGAGTGUAGC A/iBiodT/TGCUACACTCTTTCCCTACACGACGCTCTTCCG

TrAEL-seq adaptor 1 multiplexing index 5:

/5Phos/CCTTNNNNNNNNAGATCGGAAGAGCGTCGTGTAGGGAAAGAGTGUAGC A/iBiodT/TGCUACACTCTTTCCCTACACGACGCTCTTCCG

TrAEL-seq adaptor 1 multiplexing index 6:

/5Phos/GGAANNNNNNNNAGATCGGAAGAGCGTCGTGTAGGGAAAGAGTGUAGC A/iBiodT/TGCUACACTCTTTCCCTACACGACGCTCTTCCG

TrAEL-seq adaptor 1 multiplexing index 7:

/5Phos/GCACNNNNNNNNAGATCGGAAGAGCGTCGTGTAGGGAAAGAGTGUAGC A/iBiodT/TGCUACACTCTTTCCCTACACGACGCTCTTCCG

TrAEL-seq adaptor 1 multiplexing index 8:

/5Phos/TGGCNNNNNNNNAGATCGGAAGAGCGTCGTGTAGGGAAAGAGTGUAGC A/iBiodT/TGCUACACTCTTTCCCTACACGACGCTCTTCCG

where /5Phos/ indicates 5’ phosphate and /iBiodT/ indicates Biotin-dT. Adaptors were adenylated using the 5’ DNA adenylation kit (NEB, E2610S) as follows: 500 pMol DNA oligonucleotide, 5 *µ*L 10x 5’ DNA Adenylation Reaction Buffer, 5 *µ*L 1 mM ATP, 5 *µ*L Mth RNA Ligase in a total reaction volume of 50 *µ*L were incubated for 1 hr at 65 °C then 5 min at 85

°C. The reaction was extracted with phenol:chloroform pH 8, then ethanol precipitated with 10 *µ*L 3M NaOAc, 1 *µ*L GlycoBlue (Thermo AM9515), 330 *µ*L 100% (v/v) ethanol and resuspended in 50 *µ*L 0.1x TE. All adenylated adaptors were stored at -30 °C.

TrAEL-seq adaptor 2 was purchased PAGE purified from Merck or (latterly) IDT. Sequence:

/5Phos/GATCGGAAGAGCACACGTCTGAACTCCAGTCUUUUGACTGGAGTTCAGAC GTGTGCTCTTCCGATC*T

where /5Phos/ indicates 5’ phosphate and * indicates a phosphorothioate bond. Adaptor was annealed in a total reaction volume of 200 *µ*L with 20 *µ*L 100 pM/*µ*L oligonucleotide and 20 *µ*L 10x T4 DNA ligase buffer (NEB, B0202S) and heated at 95 °C for 5 min. The reaction was removed from heat and left to cool to room temperature over approximately 2 hr.

### Embedding cells in agarose

Cells were resuspended in a volume of 60 *µ*L L-buffer (100 mM EDTA pH 8, 10 mM Tris pH 7.5, 20 mM NaCl) per plug (∼1 × 10^6^ cells per plug) before distribution into 2 mL round-bottom tubes (one tube per plug). Each tube was incubated at 50 °C for 5 min, meanwhile an aliquot of CleanCut Agarose (Bio-Rad, 1703594) was heated until molten. To the 60 *µ*L cell suspension, 40 *µ*L of molten Cleancut Agarose was added and immediately vortexed for 10 s, then 100 *µ*L of solution was transferred into a disposable plug mould (Bio-Rad, 170-3713). The plug mould was placed on its side at 4 °C for 30 min before pushing plugs out into 2 mL tubes containing 500 *µ*L of fresh mammalian digestion buffer (100 mM EDTA pH 8, 10 mM Tris pH 7.5, 20 mM NaCl, 1% (v/v) N-lauroyl sarcosine, 0.1 mg/mL proteinase K).

Plugs were incubated at 50 °C overnight. Plugs were then rinsed with 1 mL TE, then washed 3 times with 1 mL TE for 1-2 hr at room temperature with rocking, and 10 *µ*L of 100 mM PMSF (Merck, 93482) was added to the second and third washes from 100 mM stock. Plugs were incubated in 200 *µ*L 1x TE containing 1 *µ*L of RNase T1 (Thermo, EN0541) for 1 hr at 37 °C. The RNase solution was then removed and plugs were washed with 1 mL 1x TE for 1 hr before storing at 4 °C in 1 mL 1x TE until use (plugs are stable for >1 yr).

### A-tailing and adaptor 1 ligation

Half of an agarose plug was used per library by cutting with a razor blade. From this point onwards the use of the term plug relates to use of a half plug. Plugs were placed in 2 mL round bottomed tubes, as with all reactions prior to agarose digestion to prevent the plug from breaking, and equilibrated once in 100 *µ*L 1 × TdT buffer (NEB, B0135S) for 30 min at room temperature, then incubated for 2 hr at 37 °C in 100 *µ*L 1 × TdT buffer containing 0.4 *µ*L 100 mM ATP (Roche, 11140965001) and 2 *µ*L Terminal Transferase (NEB, M0315L). Plugs were rinsed with 1 mL Tris buffer (10 mM Tris HCl pH 8.0), then equilibrated in 100 *µ*L 1 × T4 RNA ligase buffer (NEB, B0216S) containing 40 *µ*L 50% (w/v) PEG 8000 for 1 hr at room temperature then incubated overnight at 25 °C in 100*µ*L 1 × T4 RNA ligase buffer (NEB, B0216S) containing 40 *µ*L 50% (w/v) PEG 8000, 1 *µ*L 10 pM/*µ*L TrAEL-seq adaptor 1 with multiplex index and 1 *µ*L T4 RNA ligase 2 truncated KQ (NEB M0373L). Plugs were then rinsed with 1 mL Tris buffer, transferred to individual 15 mL tubes and washed three times in 10 mL Tris buffer with rocking at room temperature for 1-2 hr each, then washed again overnight under the same conditions.

### Agarose digestion and DNA extension

Up to 3 plugs with different multiplex indexes were transferred to 1.5 mL Eppendorf tubes and equilibrated for 15 min with 1 mL agarase buffer (10 mM Bis-Tris-HCl, 1 mM EDTA pH 6.5) before the supernatant was removed and replaced with 50 *µ*L agarase buffer per plug. Plugs were melted for 20 min at 65°C, then transferred for 5 min to a heating block pre-heated to 42 °C, followed by addition of 1 *µ*L per plug β-agarase (NEB, M0392S) and flicking to mix the contents without allowing the sample to cool. Tubes were then incubated at 42 °C for 1 hr. DNA was ethanol precipitated by adding 25 *µ*L per plug 10 M NH4OAc, 1 *µ*L per plug GlycoBlue, and 330 *µ*L per plug 100% (v/v) ethanol and vortexing, after which the pellet was washed in 70% (v/v) ethanol and then resuspended in 10 *µ*L per plug 0.1x TE (mammalian DNA samples were heated at 65 °C for 10 min to aid resuspension). DNA from up to 6 plugs with different multiplex indexes were combined in a single tube at this point.

Next 40 *µ*L per plug of reaction mix containing 5 *µ*L Isothermal amplification buffer (NEB, B0537S), 3 *µ*L 100 mM MgSO4 (NEB, B1003S), 2*µ*L 10mM dNTPs and 1*µ*L Bst 2 WarmStart DNA polymerase (NEB, M0538S) was added to the sample and incubated 30 min at 65 °C before ethanol precipitation with 12.5 *µ*L per plug 10 M NH4OAc, 1 *µ*L per plug GlycoBlue, and 160 *µ*L per plug 100% (v/v) ethanol and re-dissolving the pellet in 130 *µ*L total volume (irrespective of the number of plugs) 1x TE for 10 min at 65 °C.

### DNA shearing and streptavidin bead binding

From this point onwards the protocol is identical for 1-6 multiplexed plugs.

The DNA was transferred to an AFA microTUBE (Covaris, 520045) and fragmented in a Covaris E220 using duty factor 10, PIP 175, Cycles 200, Temp 11 °C, then transferred to a 1.5 mL tube containing 8 *µ*L pre-washed Dynabeads MyOne streptavidin C1 beads (Thermo Fisher Scientific, 65001) re-suspended in 300 *µ*L 2xTN (10 mM Tris pH 8, 2 M NaCl) along with 170 *µ*L water (total volume 600 *µ*L) and incubated 30 min at room temperature on a rotating wheel. Beads were washed once with 500 *µ*L 5 mM Tris pH 8, 0.5 mM EDTA, 1 M NaCl, 5 min on wheel and once with 500 *µ*L 0.1x TE for 5 min on wheel before re-suspension in 25 *µ*L 0.1x TE.

### Sequencing library preparation

TrAEL-seq adaptor 2 was added using a modified NEBNext Ultra II DNA kit (NEB, E7645S):

3.5 *µ*L NEBNext Ultra II End Prep buffer, 1 *µ*L 1 ng/*µ*L sonicated salmon sperm DNA (this is used as a carrier) and 1.5 *µ*L NEBNext Ultra II End Prep enzyme were added and reaction incubated 30 min at room temperature and 30 min at 65 °C. After cooling for 15 min, 1.25 *µ*L 10 pM/*µ*L TrAEL-seq adaptor 2, 0.5 *µ*L NEBNext ligation enhancer and 15 *µ*L NEBNext Ultra II ligation mix were added and incubated for 30 min at room temperature. The reaction mix was removed and beads were rinsed with 500 *µ*L wash buffer (5 mM Tris pH 8, 0.5 mM EDTA, 1 M NaCl), then washed twice with 1 mL wash buffer for 10 min on rotating wheel at room temperature and once for 10 min with 1 mL 0.1x TE. Libraries were eluted from beads with 11 *µ*L 1x TE and 1.5 *µ*L USER enzyme (NEB, M5505S) for 15 min at 37 °C, then again with 10.5 *µ*L 1x TE and 1.5 *µ*L USER enzyme (NEB, M5505S) for 15 min at 37°C, and the two eluates were combined. An initial test amplification was used to determine the optimal cycle number for each library. For this, 1.25 *µ*L library was amplified in 10 *µ*L total volume with 0.4 *µ*L each of the NEBNext Universal and any NEBNext Index primer with 5 *µ*L NEBNext Ultra II Q5 PCR master mix. Cycling program: 98 °C for 30 s, then 14-16 cycles (16 for 1-3 multiplexed libraries, 15 for 4-5, 14 for 6) of (98 °C for 10 s, 65 °C for 75 s), 65 °C for 5 min. Test PCR was cleaned with 8 *µ*L AMPure XP beads (Beckman, A63881) and eluted with 2.5 *µ*L 0.1x TE. 1 *µ*L of test PCR was examined on a Bioanalyser high sensitivity DNA chip (Agilent 5067-4626). The ideal cycle number was selected to bring the final library to final concentration of 2-4 nM, noting that the final library would be 2-3 cycles more concentrated than the test sample. 21 *µ*L of library was then amplified with 2 *µ*L each of NEBNext Universal and the chosen Index primer and 25 *µ*L NEBNext Ultra II Q5 PCR master mix using the same conditions as described above for calculated cycle number. The amplified library was cleaned with 40 *µ*L AMPure XP beads (Beckman A63881) and eluted with 26 *µ*L 0.1x TE, then 25 *µ*L of eluate was purified with 20*µ*L AMPure XP beads and eluted with 11*µ*L 0.1x TE. Final libraries were quality controlled and quantified by Bioanalyser (Agilent, 5067-4626) and KAPA qPCR (Roche, KK4835). Libraries were sequenced on an Illumina NextSeq 500 or (latterly) Element Biosciences AVITI as Single End by the Babraham Institute Genomics facility (35-40 million reads per mammalian sample).

### Data processing and mapping

Processing and mapping of all raw sequence files was carried out by the Babraham Bioinformatics facility using the updated pipeline developed by Laura Biggins - https://github.com/laurabiggins/TrAEL-seq. This pipeline includes trimming of the poly(T) tail, separation of multiplexed sets, separation into T and no-T datasets, copy number aware UMI deduplication and mapping. For deduplication, high quality mapped reads (always unique) were deduplicated by mapped position and UMI, whereas for reads with mapping quality ≤ 20 only the UMI was used for deduplication but since an 8bp UMI only has 65k combinations (smaller than the library) the first 10 bases of the unique read sequence are added to the UMI before deduplication. Human and mouse libraries were mapped to GRCh38 and GRCm38 respectively. All S. cerevisiae yeast libraries were mapped to the R64-1-1 genome with the 2-micron plasmid assembly.

### Data processing

Mapped BAM files were imported into Seqmonk (v1.48.2) and reads were trimmed down to 1 nucleotide at the 5’ end, representing the last nucleotide 5’ of the strand break. Quality filtering was set ≥20 for most analysis but this cutoff was removed for analysis of multi-copy regions. For read count comparisons within multiplexed sets, no normalisation was applied as the read count distribution within a multiplexed set should reflect the relative distribution of TrAEL-seq reads across the multiplexed samples. To compare across multiplexed sets, a total read count normalisation was applied to whole multiplexed sets, maintaining the relative distribution of read counts for samples within each multiplexed set.

RFD plots were calculated using the new SeqMonk strand bias quantitation option introduced for this purpose. Note that this option generates data from -100 to +100 which was scaled to -1 to +1 for plots for consistency with published RFD plots. Data was analysed in 20 kb windows spaced every 2 kb, split into positive and negative values and plotted on the same graph in different colours using GraphPadPrism.

Locations of Initiation Zones were determined using OKseqHMM [45]. TrAEL-seq reads were then summed in 50 kb windows spaced every 50 kb for the relevant datasets, filtering regions with aberrant counts (e.g.: line-specific CNVs and peri-centromeres), and exported as Annotated Probe Reports. These were processed using the ‘read_distance_distribution2.R’ script (https://github.com/laurabiggins/TrAEL-seq) which calculates mean read count as a function of distance from the nearest of a set of defined features (in this case the centres of Initiation Zones), as well as the 96% confidence interval and the number of regions used for the calculation. Datasets were truncated at 3 Mb as very few regions of mammalian genomes are >3 Mb from an Initiation Zone and therefore the confidence interval for these regions is very large.

## Data availability

Datasets generated in this work have been deposited at GEO accession GSE299123. The following publicly available datasets were also used:

TrAEL-seq: mouse B cells [GSE279992]; hESCs under original protocol [GSE154811]; IMR32 [GSE186122]; KM-12 and HCT116 [GSE253197]; mouse embryonic stem cells [GSE259364].

OK-seq: Mouse B cells [GSE116319].

PRO-seq: DLD-1 [GSM7009627]; hESC [GSM7431765]

## Acknowledgements

TrAEL-seq library sequencing and processing were performed by the Genomics (Geno06) and Bioinformatics (Bioinf01) teams at the Babraham Institute, which receive financial support from the Institute Core Capability Grant (BBSRC CCG). We thank Joana Neves for assistance with DLD-1 and NCI-H23 cell lines. VG reports being an employee and shareholder of Artios Pharma Ltd. HR reports patents for WO2021/028644A1, WO2016GB50517 20160229, WO2015118338 (A1), and US2022298587 (A1), all pending, as well as being an employee and shareholder of Artios Pharma Ltd and shareholder of Mission Therapeutics.

---

**Figure S1:**
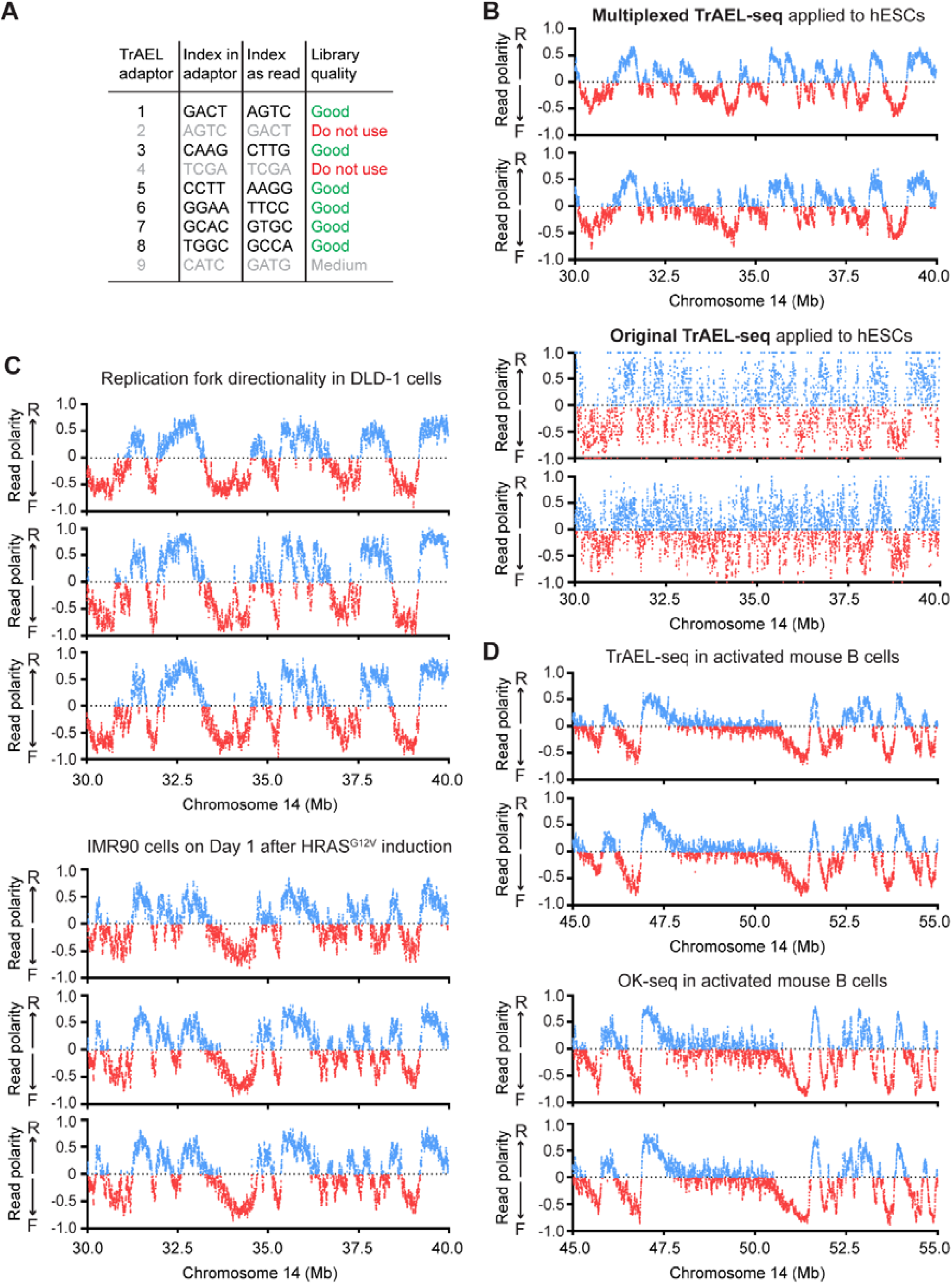
Tr AEL-seq in mammalian cells. **A**: Table of multiplexing indexes tested, the chosen set of indexes for further use was 1, 3, 5, 6, 7, 8. **B**: RFD plots of TrAEL-seq data from hESCs showing two biological replicates generated under the revised multiplexed protocol compared to two biological replicates from our original TrAEL-seq study [12]. **C**: Example RFD plots generated from 3 biological replicates each of DLD-1 cells and IMR90 cells at Day 1 of HRAS^G12V^ induction. **D**: RFD plots comparing TrAEL-seq and OK-seq data for activated mouse B cells from published data [33, 34]. All RFD plots show an arbitrary 20 Mb region, which we consider representative, the same region of the human genome is used throughout this manuscript.

**Figure S2:**
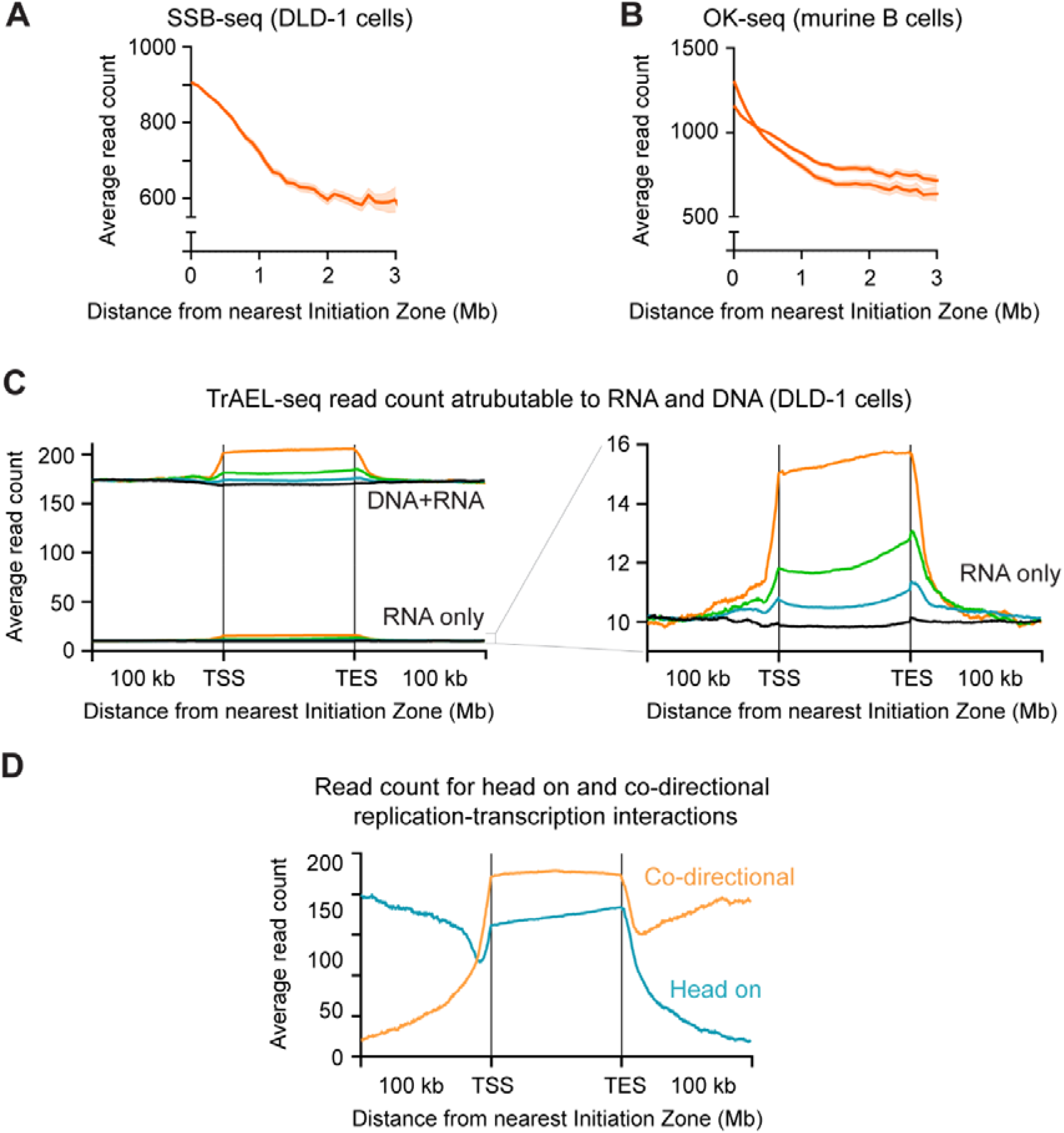
Supplement to TrAEL-seq profiling of variable replication fork density. A: Plot of total SSB-seq read count at increasing distance from replication Initiation Zones in DLD-1 cells. Solid line shows mean value, shaded band 95% confidence interval. Read counts were summed in 50kb windows spaced every 50kb, with regions of altered copy number or aberrant read count removed. Distance was calculated from Initiation Zones determined using OKSeqHMM [45], and average read count determined in 100kb windows of distance from the centre of initation zones. SSB-seq detects will detect both leading and lagging strand of replication forks as it is performed using an *in vitro* primer extension on total DNA with biotin-labelled dNTPs. B: Plot of total OK-seq read count from B cells using published data [33], solid lines show mean value, shaded band 95% confidence interval, analysed as in A. C: Metaplot of TrAEL-seq read count in DLD-1 cells averaged across genes ±100 kb as in Figure 2C, showing both reads starting with T (which could derive from DNA or RNA) and separately the reads starting with other nucleotides (which are almost entirely RNA). The RNA count has been divided by 3 to compensate for this representing reads starting with 3 of the 4 nucleotides, the point of this comparison being to indicate the proportion of the reads in the DNA+RNA library that are likely to derive from RNA; even omitting this compensation, the proportion of the DNA+RNA signal arising through TrAEL-seq detection of nascent RNA remains a small part of the signal and cannot explain the observed enrichment of TrAEL-seq reads over transcribed regions. D: Metaplots as in Figure 2C, divided into reads representing replication forks moving co-direction or head-on to transcribed regions.

**Supplementary File 1: Full TrAEL-seq protocol with trouble-shooting guide**

