## Supplementary File 1 for "TrAEL-seq captures DNA replication dynamics in mammalian cells"

**Multiplexed TrAEL-seq**

The multiplexed protocol and adaptors work so well we rarely if ever use the original protocol now.

Reagents and consumables are detailed after the protocol

Common problems and solutions are given at the end of this document

*Input material*

Start with agarose plugs containing 1-4x10^7 yeast cells that were ethanol fixed at harvest or 1-2x10^6 mammalian cells^[1]^ (see protocol for Agarose Embedding), you need ½ of a plug per sample.

*Stopping points*

This protocol is very long but can be done in bits, and we have stopped at many points without problems. To our knowledge all the following stopping points are fine:

You can pause indefinitely at any point where plugs are in tris buffer, store at 4°

You can pause indefinitely when DNA is in solution in TE, store at 4°.

You can pause indefinitely when DNA solutions have ethanol added for precipitation, is a pellet in the 70% ethanol, or the pellet is re-suspending in TE. Store at 4°.

You can pause indefinitely when DNA is bound to magnetic beads once beads are in bead wash or TE, store at 4°. Vortex well to re-suspend when you re-start.

Once you start the library preps using the NEBNext Ultra II kit, you must continue until the beads are in the wash solution, but then you can store at 4°.

After eluting the DNA from the magnetic beads with USER, store the libraries in the freezer until ready to amplify. Similarly, amplified libraries before or after purification are stored in the freezer.

The latest version of this protocol should be available from our lab website:

<https://www.babraham.ac.uk/our-research/epigenetics/jon-houseley/protocols>

*Protocol*

NOTE: At the start you are handling each sample separately, they are pooled later

Transfer ½ plug 2ml tube

Equilibrate plug once in 100µl 1x TdT buffer, 30 min at room temperature (RT)

Exchange for 100ul containing:

10ul 10x Terminal transferase buffer (NEB)

0.4ul 100mM ATP^[2]^

1ul Terminal transferase (NEB)

84ul water

Incubate 2-3 hrs at 37

Discard the solution and rinse plug with 1ml tris buffer

Then equilibrate plug 1 hr at RT with:

10ul 10x T4 RNA ligase buffer (NEB)

40ul 50% PEG 8000^[3]^

50ul water

Exchange for 100ul containing:

10ul 10x T4 RNA ligase buffer^[4]^

40ul 50% PEG 8000

1ul App TrAEL indexed adaptor (indexes 1,3,5,6,7 or 8)

48ul water

1ul T4 RNA ligase 2 KQ

Mix all components except ligase in a 1.5ml tube, vortex vigorously, then add ligase and pipette repeatedly with a P200 cut tip to mix it really well

Incubate overnight at 25deg [NOT LONGER!!!]

Discard reaction mix, rinse plug 1x 1ml tris buffer, transfer to 15ml tube

During the day wash 4x with 10ml tris buffer at room temperature on rocker

Leave overnight on rocker with 10ml tris buffer

Equilibrate 15min or more with 1ml agarase buffer^[5]^

Meanwhile, heat a block to 65° and another to 42°C ^[6]^

Now pool the libraries:

Transfer all the plugs to a single 1.5 ml tube (up to 6 plugs with different indexes)^[7]^

This is a bit fiddly. The easiest way we have found is to pour the 1ml agarase buffer + plug from the 15ml falcon tube into an intermediate 1.5ml tube and remove the buffer so that you have the plug by itself. Then, invert this over the tube in which you collecting the plugs and flick gently until it falls down into the tube with the other plugs

Remove any residual agarase buffer, add agarase buffer if necessary to a final volume of ~300 μl and melt 20 min at 65°

Shift tubes to 42° for 5 min, add 1ul β-agarase per plug, flick, incubate 1 hour at 42°

Be quick during addition – do not let the tube cool down

Add 75 µl 10M NH_4_OAc, 3 µl glycoblue and 1ml of ethanol, vortex vigorously (particularly for mammalian samples)

Chill 15 min on ice then spin 15 min top speed 4° ^[8]^

Wash with 70% ethanol, dry 5 min, resuspend in 10µl 0.1xTE per plug, leave >10 min at room temperature

For mammalian samples 15 min at 65° is better but make sure it doesn’t go dry

Per plug, add master mix containing	29µl water

5µl Isothermal amp buffer (NEB)

3µl 100mM MgSO4

2µl dNTPs (10mM)

1µl Bst 2 warmstart pol^[9]^ (NEB)

Pipette to mix and re-suspend pellet

Heat 30min at 65°

Per plug, add 12.5µl 10M NH4OAc, 1µl glycoblue and 160µl of ethanol, vortex well

Chill 15 min on ice then spin 15 min top speed 4°

Wash with 1 ml 70% ethanol, dry 5 min, resuspend in 130µl 1xTE 15min at 65°. Note that is 130 μl in total, not per plug.

Can proceed directly or store at 4°

**Note**: the following sonication protocol is for our Covaries E220, and should yield libraries of 500-600bp final size (including adaptors). If using a different sonicator, you will need to select a sonication scheme that gives average fragment sizes of 400-500bp (which will increase to 500-600bp once all adaptors are added). If you sonicate the library into smaller fragments it is harder to separate the library from the adaptor dimers. The sonication volumes are not too important as the next step is binding to streptavidin beads in 600μl of 1x TN. We don’t recommend enzymatic fragmentation methods as the ones we have tried chew up the ssDNA bits on the TrAEL-seq adaptor.

Transfer to Covaris tubes (AFA Snap-cap microTUBE)

Machine use:

Turn on machine

Turn off and restart!

Turn on computer

Turn on chiller

Fill water tank to 6L

Fill in booking sheet

Protocol (Cris in JHGroup folder)

Duty factor 10%

Peak Incident Power 175W

Cycles 200

Temp 11ºC

Treatment time 50s

Sample volume 130µl

After use turn everything off, empty water and dry

NOTE: From this point one sample = one multiplexed pool (which comes from 1-6 original samples). Do not increase reagent volume for the number of samples in the multiplexed set.

Wash 8µl Dynabeads MyOne streptavidin C1 beads per sample twice with 0.5ml bead prep buffer (5mM Tris pH8, 0.5mM EDTA, 1M NaCl, 0.05% Tween), 3 min on wheel per wash [you can wash beads for up to 4 samples together]

Resuspend beads in 300µl 2xTN (10 mM Tris pH 8, 2 M NaCl) per sample, split into 300µl aliquots in 1.5ml tubes and add 170µl water to each

Add 130µl sonicated DNA to beads^[10]^

Incubate on wheel 30 min at RT

Spin tubes up to 1000rpm before putting on magnet to collect liquid

Wash beads 1x 500µl 5mM Tris ph8, 0.5mM EDTA, 1M NaCl, 5 min on wheel

Spin tubes up to 1000rpm before putting on magnet to collect liquid

Wash beads 1x with 500µl 0.1x TE, 5min on wheel

Spin tubes up to 1000rpm before putting on magnet to collect liquid

Carefully remove the wash solution so as not to leave any drops and re-suspend beads immediately in 25 µl 0.1x TE

Store overnight at 4° or carry on directly

Add	3.5μl NEBNext Ultra II end repair buffer

1µl 1ng/ul salmon sperm DNA^[11]^

1.5μl NEBNext Ultra II end repair enzyme

Pipette to mix and incubate 30 min at 20º (room temperature) then 30 min at 65º

Let cool then spin 10s at 1000rpm

Add	1.25μl new ENDseq adaptor 2 (from annealed 10uM stock)^[12]^

0.5μl ligation enhancer

15μl NEBNext Ultra II ligation mix

Pipette to mix and incubate 30 min at 20º (room temperature)

Remove reaction mix using magnet, rinse beads 500ul wash buffer (5mM Tris ph8, 0.5mM EDTA, 1M NaCl). REMEMBER your libraries are on the beads!

Wash beads 2x 1ml wash buffer (5mM Tris ph8, 0.5mM EDTA, 1M NaCl) 10 min on wheel each

Spin tubes up to 1000rpm before putting on magnet to collect liquid

Wash beads 1x with 1ml 0.1x TE, 10min on wheel

Spin tubes up to 1000rpm before putting on magnet to collect liquid

Carefully remove the wash solution so as not to leave any drops and re-suspend beads immediately in 11µl 1x TE + 1.5μl USER enzyme

Incubate 15 min at 37º

This elutes the DNA from the beads

Collect beads using magnet, transfer 12µl supernatant to a 1.5ml LoBind tube using an orange LoBind tip

The supernatant contains the library but we do another round of elution to get anything left. This is the first step in which the very low concentration DNA is unprotected, hence the LoBind tubes and tips

Resuspend beads in 10.5µl 1x TE + 1.5μl USER enzyme

Incubate 15 min at 37º

This elutes any remaining DNA from the beads

Collect beads using magnet, add 12µl supernatant to the 12µl from the first elution using an orange LoBind tip

Store samples at -20 if necessary

We then perform a test amplification to work out how many cycles we need to use for the real library.

In PCR tubes, mix	1.25μl library

0.4μl NEBNext any index oligo

0.4μl NEBNext universal oligo

2.95ul water

5μl NEBNext Ultra II PCR mix

98º 30s

98º 10s \

65º 75s / x 14 cycles^[13]^

65º 5 min

4º hold

Clean with 8μl AMPure beads, elute in 2.5μl 0.1x TE

You can elute up to 2.0μl from these but it is easier just to put the eluate and bead mix on the magnet and pipette 1µl straight onto the Bioanalyzer lane

Run on Bioanalyzer

Use this to calculate the cycle number for the final amplification – note: because of the various dilutions, assume that the final library would be 4x more concentrated than the test library, so reduce the number of cycles by 2 before further adjustments. Try to aim for a final library concentration of 1.5-4 nM. Adaptor dimers are often seen in these test reactions, be sure to exclude from the quantification.

You should quantify across the region from 300-1,000 bp, which should exclude the adaptor peaks. However, it is not uncommon to get weird profiles like these:

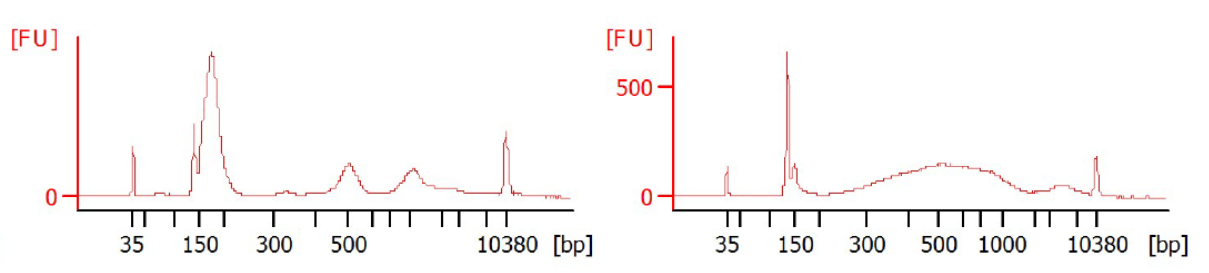

Don’t worry – your libraries should be fine, but quantification is not so accurate. Err on the side of slightly over amplifying the library (so only reduce the number of cycles by 1 rather than 2 before adjusting the concentration). Library over-amplification doesn’t seem to cause big problems in TrAEL-seq.

Final library amplification:

In PCR tubes, mix	21μl library

2μl NEBNext index oligo

2μl NEBNext universal oligo

25μl NEBNext Ultra II PCR mix

98º 30s

98º 10s \

65º 75s / x calculated cycles

65º 5 min

4º hold

Transfer libraries to 1.5 ml tubes for final purification (purification seems to be better in 1.5 ml tubes than in PCR tubes), and make sure they are still 50μl (top up with water if not).

Clean with 40μl AMPure beads, elute with 26μl 0.1xTE for 25μl yield

Clean again with 20μl AMPure beads, elute with 11μl 0.1xTE for 10.5μl yield

Run 1µl directly on Bioanalyser and use 0.2µl for KAPA quantification [see separate protocol]

It is not unusual to have some residual adaptor in the library. If so, dilute to 25ul and perform another 2 rounds of AMPure bead purification at 0.8x.

**Important sequencing info**:

**It is critical that at least 10% of some library that is not multiplexed TrAEL-seq be included in the sequencing lane!**

This is because the invariant T read as the first base after the barcode crashes the base calling algorithm (at least on our NextSeq 500). This causes the surrounding bases in the barcode to be mis-read and not assigned. Spiking in another type of library fixes it.

**Barcodes for TrAEL-seq index adaptors:**

We currently have a working set of 6 adaptors - indexes 1, 3, 5, 6, 7, 8

|  | In the adaptor | What is actually read | Quality |
| --- | --- | --- | --- |
| **1** | **GACT** | **agtc** | **Good** |
| ~~2~~ | ~~AGTC~~ | ~~gact~~ | ~~Poor - Not in use~~ |
| **3** | **CAAG** | **cttg** | **Good** |
| ~~4~~ | ~~TCGA~~ | ~~tcga~~ | ~~Poor - not in use~~ |
| **5** | **CCTT** | **aagg** | **Good** |
| **6** | **GGAA** | **ttcc** | **Good** |
| **7** | **GCAC** | **gtgc** | **Good** |
| **8** | **TGGC** | **gcca** | **Good** |
| **~~9~~** | **~~CATC~~** | **~~gatg~~** | **~~Maybe poor - not in use?~~** |

As an example this is the structure of TrAEL index 1

**TrAEL index 1**

/5Phos/**GACT**NNNNNNNNAGATCGGAAGAGCGTCGTGTAGGGAAAGAGTGUAGCA/iBiodT/TGCUACACTCTTTCCCTACACGACGCTCTTCCG

Note: the quality of these adaptors synthesised by Sigma-Genosys was poor. We have had all of them synthesised by IDT with very good results.

**Reagents**

*Adaptors*

TrAEL-seq index 1:

/5Phos/**GACT**NNNNNNNNAGATCGGAAGAGCGTCGTGTAGGGAAAGAGTGUAGCA/iBiodT/TGCUACACTCTTTCCCTACACGACGCTCTTCCG

TrAEL-seq index 3:

/5Phos/**CAAG**NNNNNNNNAGATCGGAAGAGCGTCGTGTAGGGAAAGAGTGUAGCA/iBiodT/TGCUACACTCTTTCCCTACACGACGCTCTTCCG

TrAEL-seq index 5:

/5Phos/**CCTT**NNNNNNNNAGATCGGAAGAGCGTCGTGTAGGGAAAGAGTGUAGCA/iBiodT/TGCUACACTCTTTCCCTACACGACGCTCTTCCG

TrAEL-seq index 6:

/5Phos/**GGAA**NNNNNNNNAGATCGGAAGAGCGTCGTGTAGGGAAAGAGTGUAGCA/iBiodT/TGCUACACTCTTTCCCTACACGACGCTCTTCCG

TrAEL-seq index 7:

/5Phos/**GCAC**NNNNNNNNAGATCGGAAGAGCGTCGTGTAGGGAAAGAGTGUAGCA/iBiodT/TGCUACACTCTTTCCCTACACGACGCTCTTCCG

TrAEL-seq index 8:

/5Phos/**TGGC**NNNNNNNNAGATCGGAAGAGCGTCGTGTAGGGAAAGAGTGUAGCA/iBiodT/TGCUACACTCTTTCCCTACACGACGCTCTTCCG

Purchase PAGE purified from IDT

These must be 5’ adenylated (5’ App) before use using the 5’ adenylation kit (see separate protocol for this)

TrAEL-seq adaptor 2 (aka: ENDseq adaptor 2):

[Phos]GATCGGAAGAGCACACGTCTGAACTCCAGTCUUUUGACTGGAGTTCAGACGTGTGCTCTTCCGATC*T

Purchase PAGE purified from IDT

Annealing:

20µl 100uM ENDseq Adaptor 2 in 200ul 1x T4 DNA ligase buffer

Incubate in heating block at 95°C 5 min then remove block from the heat and leave to cool to room temperature

*Reagents and consumables*

Terminal Transferase (NEB M0315S)

100mM ATP (Merck) aliquot as 5ul in PCR tubes on receipt and store at -80°

T4 RNA ligase 2 truncated KQ (NEB M0373L)

T4 RNA ligase buffer (NEB supplied with T4 RNA ligase 2)

50% PEG 8000 (NEB supplied with T4 RNA ligase 2 or home-made as required)

β-agarase (NEB M0392S)

GlycoBlue (Thermo AM9515)

1µl Bst 2 WarmStart DNA polymerase (NEB M0538S)

Isothermal amplification buffer (NEB supplied with Bst pol)

100 mM MgSO_4_ (NEB supplied with Bst pol)

10 mM dNTPs

Ethanol

AFA microTUBEs (Covaris 520045)

Dynabeads MyOne streptavidin C1 beads (Thermo, 65001)

NEBNext Ultra II DNA kit (NEB E7645S), additional Q5 mix may be required which can be purchased separately from NEB

1ng/µl sonicated salmon sperm DNA or other carrier

NEBNext Multiplex Oligos set (e.g. NEB E7335S), additional USER mix may be required which can be purchased separately from NEB

AMPure XP beads (Beckman A63881)

Bioanalyser high sensitivity DNA chip (Agilent 5067-4626)

KAPA qPCR (Roche KK4835) or equivalent library quantification kit

*Solutions*

These solutions are made with milliQ water

Tris buffer	10mM Tris pH8.0

Agarase buffer	10 mM Bis-Tris-HCl, 1 mM EDTA pH 6.5

These solutions are made with certified DNase/RNase free water

10 M NH_4_OAc

1x TE and 0.1x TE	Diluted from 100x stock

2xTN	10 mM Tris pH 8, 2 M NaCl

Wash buffer	5 mM Tris pH 8, 0.5 mM EDTA, 1 M NaCl

Bead prep buffer	5mM Tris pH8, 0.5mM EDTA, 1M NaCl, 0.05% Tween

*Equipment*

Magnetic rack for 1.5 ml tubes

Magnetic rack for 0.2 ml tubes (optional)

Covaris E220 sonicator

Other sonicators may be suitable, protocol adjustments will be required

PCR machine

Agilent Bioanalyzer or equivalent gel system

Wheel

Rocker

2 heating blocks

LoBind 1.5ml tubes

LoBind tips for P20

**Common TrAEL-seq problems and solutions**

*Cell number*

For yeast, samples of <1x10^6^ cells perform poorly, 1x10^7^ seems optimal. For mammalian cells 1x10^6^ is optimal, less than 1x10^5^ or more than 1x10^7^ cells can impair the outcome. Note that since half an agarose plug is used for the library you should double these numbers for the amount of cells to embed in a plug.

*Adaptor synthesis*

We and others have found some oligonucleotide manufacturers produce poor quality adaptors or fail entirely. We now use IDT as a supplier for these demanding oligonucleotides, and results have so far been excellent, particularly with the redesigned sequences presented here.

*Sequencing*

TrAEL-seq libraries should never occupy more than 90% of a sequencing lane as the invariant T’s at the ligation junction cause base-calling errors for surrounding nucleotides. Data can be rescued by trimming 3nt from the start of the dead, but assigning indexes becomes very difficult.

*Fragmentation*

TrAEL-seq libraries should be fragmented to fairly large fragments such that the final average library size is 600-800nt. Much smaller and the libraries are hard to separate from the often large amount of adaptor dimers produced. Other sonication systems should be effective, but some chemical fragmentation systems will cleave single stranded regions of TrAEL-seq adaptors.

*Test amplification*

This can often yield strange bioanalyser profiles. Don’t worry, final libraries are usually fine and quantitation based on the 200-1000bp region remains fairly accurate.

*Excessive adaptor diners*

Although most libraries amplify well, some have problems particularly if the input amount is low. We have noted that performing a 0.8x AMPure cleanup after USER elution prior to PCR can dramatically reduce adaptor dimers.

*Yield and purity*

We originally suggested aiming for 1nM final library concentration, but experience shows this to be a bit low and we now aim for 2-4nM. This allows libraries to be further AMPure cleaned at 0.8x to remove residual adaptor dimers (not uncommon). Additional PCR cycles can be performed if final library concentration is low, but a further two rounds of AMPure cleanup are required afterwards.

*Choice of multiplex sets*

Differences in TrAEL-seq substrate amounts will lead to differences in the read output across multiplexed sets. Some cell lines give particularly strong TrAEL-seq signals (we have seen this with SW620 for example), and will tend to swamp other samples multiplexed in the same pool such that the majority of reads stem from a few libraries while others are under-sequenced. We therefore recommend not mixing cell lines within a pool, and if detailed analysis of a poorly replicating cell type is required (eg: some drug treated samples), it is wise not to mix these with untreated controls.

1. Ethanol fixing of mammalian cells yet to be tested
2. Stored in 5ul aliquots in the -80 from a Merck stock solution
3. Use home-made 50% PEG – the kit does not come with enough
4. Care! Not T4 DNA ligase buffer!
5. Home made, but same as NEB agarase buffer
6. Make sure these really reach temperature before use! Give them at least 20 min
7. This is a recent modification. Almost all multiplexed libraries have been made by pooling up to 3 plugs in a tube and adding 50ul per plug of agarase buffer. Two tubes, each containing 3 plugs, have then been combined after precipitation but before the Bst step. This modification does not seem to make a difference and is simpler.
8. Sample didn’t pellet and formed a weird gelatinous lump? We’ve seen this with HeLa samples. We added some 3M NaOAc and some more ethanol and re-span, that sorted the problem.
9. Don’t use beyond the expiry date – you get lots in a tube but this enzyme can go off and that causes subtle but significant problems
10. With our current filter tips it is very hard to remove material from Covaris tubes – use a normal yellow tip instead
11. This is included to buffer the voracious activity of T4 DNA polymerase in the End repair mix so it doesn’t degrade rare DNA fragments. It may not be necessary.
12. Note – do NOT use the ENDseq 2c adaptor – it will not work!
13. This is for 4-6 multiplexed mammalian samples, use 15 cycles for 2-3 samples or 16 cycles for 1 sample. Increase by 1 for yeast samples or if you have reason to believe your signals will be low. It is not critical you get this right, you can always do another test PCR if your amplification is wildly under or over. The whole point is to be able to calculate the perfect cycle number for the final amplification.
